# New insights on the modelling of the molecular mechanisms underlying neural maps alignment in the midbrain

**DOI:** 10.1101/2020.06.15.152777

**Authors:** Elise Laura Savier, James Dunbar, Kyle Cheung, Michael Reber

## Abstract

We previously identified and modelled a principle of visual maps alignment in the midbrain involving the mapping of the retinal projections and concurrent transposition of retinal guidance cues into the superior colliculus providing positional information for the organization of cortical V1 projections onto the retinal map (Savier et al., 2017). This principle relies on mechanisms involving Epha/Efna signaling, correlated neuronal activity and axon competition. Here, using the 3-step map alignment computational model, we predict and validate in vivo the visual mapping defects in a well-characterized mouse model. Our results challenge previous hypotheses and provide an alternative, although complementary, explanation for the phenotype observed. In addition, we propose a new quantification method to assess the degree of alignment and organization between maps, allowing inter-model comparisons. This work generalizes the validity and robustness of the 3-step map alignment algorithm as a predictive tool and confirms the basic mechanisms of visual maps organization.

## Introduction

Understanding and modelling the mechanisms of neural circuits formation in the brain has been a challenging subject in fundamental neurobiology for decades. One of the most studied biological models to investigate the formation and function of sensory connectivity -or sensory maps-from both experimental and theoretical standpoints is the superior colliculus (SC), an evolutionary conserved structure located in the midbrain (Cang and Feldheim, 2013; Basso and May, 2017). In most vertebrates, the SC is the premier brain center for integrating sensory inputs from visual, auditory and somatosensory modalities distributed in different interacting laminae. This structure is a key node in the network of brain areas responsible for controlling the location of attention and even decision-making (Basso and May, 2017). Visual inputs in the superficial layers of the SC correspond to the organized projections from the retinal ganglion cells (RGCs) in the retina (the retino-collicular projections) and from layer V neurons in the primary visual cortex V1 (the cortico-collicular projections). Both projections form visual maps that must be aligned and in register to allow efficient detection of visual stimuli (Basso and May, 2017; Liang et al., 2015; Zhao et al., 2014).

Although many studies have focused on the mechanisms involved in the formation of the retino-collicular map, little is known about how the retino- and cortico-collicular maps are aligned during development. Previous studies on retino-collicular mapping have revealed the involvement of gradients of Eph tyrosine kinases receptors (Eph) and their cognate membrane-bound ligands, the ephrins (Efn), together with competition between axons for collicular space and correlated neuronal activity in the form of spontaneous waves (Ackman and Crair, 2014; Cang and Feldheim, 2013; Triplett et al., 2011). To gain insight into the molecular mechanisms involved in map formation, mouse genetic models were generated, in which the expression of members of the Eph and ephrin family was manipulated. The mapping mechanisms of the nasal-temporal axis of the retina onto the rostral-caudal axis of the SC involving Epha/Efna signaling have been, by far, the most studied experimentally (Feldheim et al., 1998; Brown et al., 2000; Feldheim et al., 2000; Reber et al., 2004; Rashid et al., 2005; Pfeiffenberger et al., 2006; Triplett et al., 2009; Bevins et al., 2011; Suetterlin and Drescher, 2014; Owens et al., 2015; Savier et al., 2017; Savier and Reber, 2018). Several hypotheses have been made to account for the phenotypes observed in mouse genetic models and this extensive body of work has been used to generate computational approaches which attempt to replicate experimental findings (Honda, 2003; Reber et al., 2004; Honda, 2004; Koulakov and Tsigankov, 2004; Goodhill and Xu, 2005; Willshaw, 2006; Tsigankov and Koulakov, 2006, 2010; Triplett et al., 2011; Simpson and Goodhill, 2011; Grimbert and Cang, 2012; Sterratt and Hjorth, 2013; Willshaw et al., 2014; Hjorth et al., 2015; Owens et al., 2015; Savier et al., 2017). However, until recently, these models have not been able to explain how collicular visual maps are aligned during development.

Interestingly, the formation of the retino-collicular map occurs prior to the establishment of the cortico-collicular map. Other studies have shown that the existence of the retino-collicular map is necessary for the formation of the cortico-collicular map, suggesting an interdependence of the two mechanisms (Khachab and Bruce, 1999; Rhoades et al., 1985; Triplett et al., 2009). Similar observations have been made in other part of the visual system (Shanks et al., 2016). Another piece of evidence came from the study of map alignment in the Isl2-Epha3KI, one of the best characterized mutant in the field. In these mutants, the Epha3 receptor is ectopically expressed in 50% of RGCs, leading to a duplication of the retino-collicular map (Brown et al., 2000). Strikingly, a full duplication of the cortico-collicular map is also observed in the Isl2-Epha3^KI/KI^ homozygous mutants, which display a normal retinotopy in the visual cortex (Triplett et al., 2009). The authors concluded that a retinal-matching mechanism involving spontaneous correlated activity in the retina instructs cortico-collicular projections and alignment onto the retino-collicular map. Alternative explanations can also be suggested. For instance, the ectopic expression of Epha3 specifically in the retina may alter the expression of other members of Epha/Efna throughout the visual system disrupting maps formation and alignment. In another example, molecular cues originating from the retina could be carried over to the colliculus and provide mapping/alignment information to ingrowing cortical axons, as suggested earlier (Savier et al., 2017). This latter hypothesis is in line with a retinal-matching mechanism inferred by Triplett and collaborators (Triplett et al., 2009). A similar mechanism of guidance cues transportation has recently been demonstrated for axon guidance at the optic chiasm in the mouse visual system (Peng et al., 2018).

To gain insight into the implication of molecular guidance cues in the retina in the alignment of visual maps in the SC, we have characterized a new mutant, the Isl2-Efna3KI, which over-expresses the ligand Efna3 in 50% of the RGCs (Savier et al., 2017). To our surprise, this mutant does not display any defect in the formation of the retino-collicular map, however the subsequent cortico-collicular map is duplicated. This led us to conclude that molecular guidance cues expressed in the retina are implicated in the formation cortico-collicular map (Savier et al., 2017). To simulate this mechanism *in silico*, we generated the 3-step map alignment model, based on the Koulakov model (Koulakov and Tsigankov, 2004; Tsigankov and Koulakov, 2010, 2006). Many different algorithms have been generated to model retino-collicular mapping controlled by Eph/Efn. Recently, Hjorth et al., (Hjorth et al., 2015) developed a pipeline which allowed for systematic testing of the currently available models and revealed that most cannot reproduce all nuances of experimental findings (Isl2-Epha3KI, EfnaKOs and Math5KO). Among those tested, the most faithful was the Koulakov model, which had recently been extended to explain the variability in the phenotypes observed in a particular mouse model, the Isl2-Epha3KI (Owens et al., 2015).

The 3-step map alignment model simulates the formation of the retino-collicular map and, for the first time, the subsequent formation and alignment of the cortico-collicular map along the rostral-caudal axis of the SC based on experimental and mechanistic evidence (Savier et al., 2017; Savier and Reber, 2018). This algorithm predicts normal wildtype (WT) mapping as well as the map alignment defects observed in the Isl2-Efna3KI animals (Savier et al., 2017). However, whether it also simulates mapping abnormalities observed in other Epha/Efna mutants, and particularly in the Isl2-Epha3KI animal model, is unknown. Here we demonstrate that the 3-step map alignment algorithm accurately simulates both retino- and cortico-collicular mapping defects in Isl2-Epha3KI mutants (Brown et al., 2000; Triplett et al., 2009). Our results strongly suggest that the mechanism underlying the subsequent duplication of the cortico-collicular projection corresponds to a redistribution of retinal molecular cues, the Efnas, into the SC. The retinal Efnas, provided by the incoming retinal axons within the SC, act together with correlated activity to instruct cortico-collicular alignment onto the retino-collicular map (Savier et al., 2017; Triplett et al., 2009). We further confirmed and validated the predictions of the algorithm by quantitative *in vivo* map analysis in both heterozygous and homozygous Isl2-Epha3KI animals. Moreover, a new implementation of the algorithm generates indexes providing a qualitative measure of map organization and allowing comparison of visual map layouts between biological models. Together with our previous work (Savier et al., 2017), these data confirm the validity and robustness of our algorithm and reinforces the underlying principle of visual maps formation and alignment in the midbrain, where the layout of the dominant retino-collicular map specifies the alignment of the cortico-collicular map through spontaneous correlated activity and transposed retinal Efnas. Such principle dictates the optimal functioning of the system by allowing fine adjustments which compensate for intrinsic variability - or stochasticity-of sensory maps formation.

## Results

### Normal Ephas/Efnas expression in Isl2-Epha3^KI/KI^ retinas, SC and V1 cortex

Retinal expression of *Epha* receptors in Isl2-Epha3KI animals has been extensively characterized (Reber et al., 2004). However, whether EPHA3 ectopic expression in Isl2-positive (Isl2(+)) RGCs affects retinal *Efna* co-expression is unknown. We performed a quantitative analysis of retinal *Efna* transcripts in P1/P2 nasal, central and temporal acutely isolated RGCs (Savier et al., 2017), revealing similar expression levels between Isl2-Epha3^KI/KI^ and WT littermates. The graded expression of *Efna2* and *Efna5* running form high-nasal to low-temporal is preserved in Isl2-Epha3^KI/KI^ animals, whereas the expression of *Efna3* is homogeneous (Fig 1A). Transcript analyses in V1 and SC (Fig 1B) revealed similar levels of *Efna2/a3/a5* and *Epha4/a7* receptors in Isl2-Epha3^KI/KI^ compared to WT littermates at P7 excluding indirect effects on mapping due to local changes of *Ephas/Efnas* gene expression.

**Figure 1.**
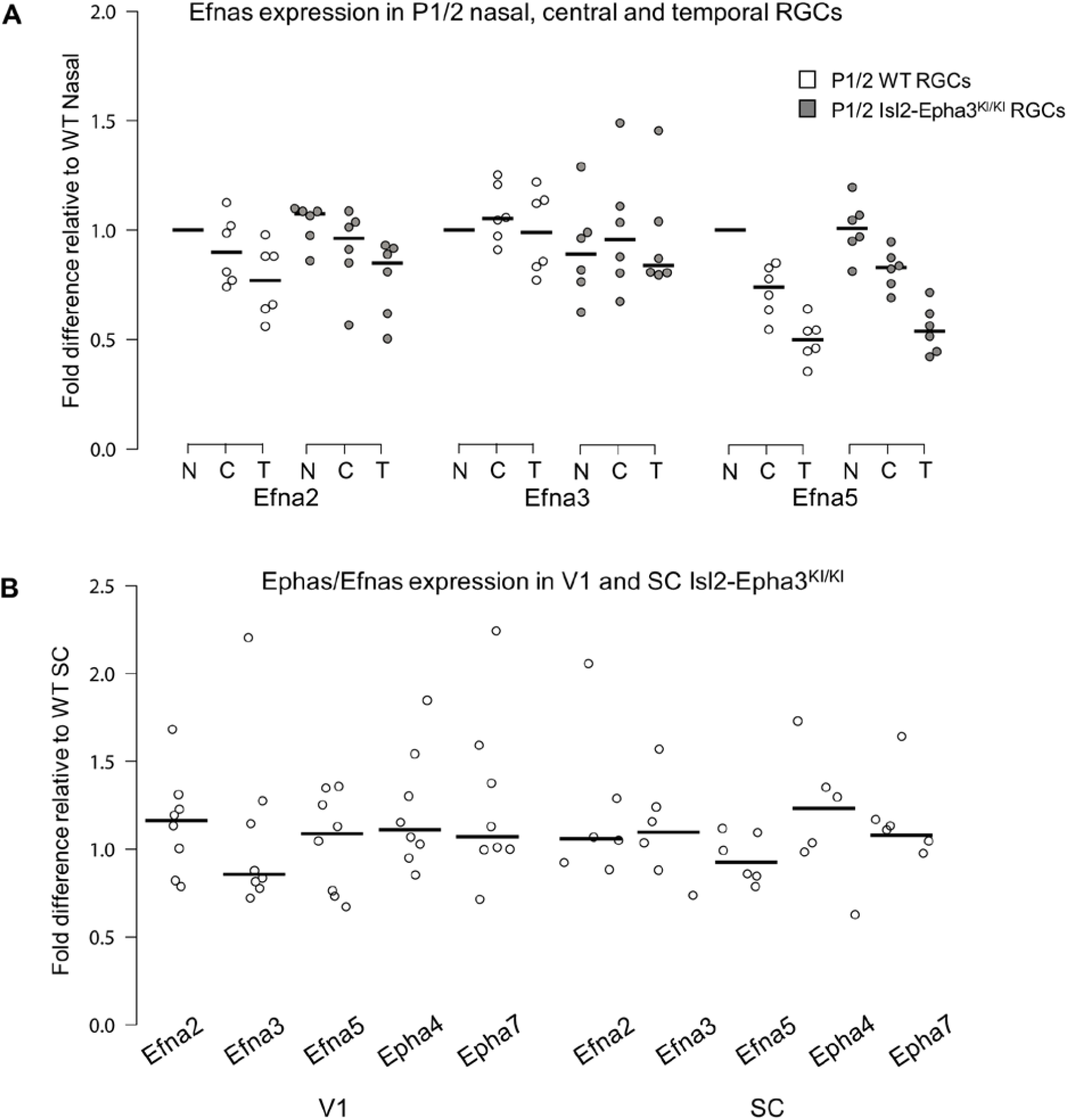
Dot plots representing Ephas/Efnas expression in Isl2-Epha3^KI/KI^ retinas, V1 cortex and SC. **A.** Median Efna2/a3/a5 expression levels (relative to wildtype nasal expression) in P1/2 wildtype (WT - white) and Isl2-Epha3^KI/KI^ (grey) acutely isolated RGCs from nasal (N), central (C) and temporal (T) retinas (WT, Isl2-Epha3^KI/KI^, n = 6 animals, 12 retinas, Two-way ANOVA without replication: Efna2 x genotype: F_(1,2)_ = 3.72 < F_crit._ = 18.5, p = 0.19; Efna3 x genotype : F_(1, 2)_ = 11.13 < F_crit._ = 18.5, p = 0.07; Efna5 x genotype: F_(1, 2)_ = 3.58 < F_crit._ = 18.5, p = 0.20 **B.** Median Efna2/a3/a5 ligands and Epha4/a7 receptors expression levels (relative to WT expression levels) in Isl2-Epha3^KI/KI^ V1 (WT n = 5 animals, Isl2-Epha3^KI/KI^ n = 8 animals; variables are normally distributed, one sample t-test: Efna2: p = 0.29; Efna3: p = 0.43; Efna5: p = 0.42; Epha4: p = 0.07; Epha7: p = 0.54) and SC (WT n = 5 animals, Isl2-Epha3^KI/KI^ n = 6 animals; variables are normally distributed, one sample t-test: Efna2: p = 0.20; Efna3: p = 0.65; Efna5: p = 0.71; Epha4: p = 0.11; Epha7: p = 0.17). qPCRs were repeated 3 times with duplicates for each sample.

### Simulation of the Isl2-Epha3KI mutants retino- and cortico-collicular mapping

Our data above, together with previous work (Reber et al., 2004; Savier et al., 2017), showed that ectopic expression of Epha3 or Efna3 in Isl2(+) RGCs does not affect endogenous Efnas nor Ephas expression within the retina, SC and V1. We can reasonably assume that the equations modelling retinal, collicular and cortical Efnas gradients as well as cortical Ephas gradients related to the Isl2-Epha3KI genotypes are equivalent to the equations previously characterized (measured experimentally or estimated) in WT animals (Reber et al., 2004; Cang et al., 2005; Tsigankov and Koulakov, 2006, 2010; Bevins et al., 2011; Savier et al., 2017). Measured gradients of Epha receptors (R _Epha_) in RGCs along the nasal-temporal axis (x), R _Epha_ (x)^retina^ (Fig 2A, G, M) (Brown et al., 2000; Reber et al., 2004) are modelled by two equations, one corresponding to Isl2(+) RGCs expressing WT levels of Ephas + Epha3 (in Fig 2G, M) and the second corresponding to Isl2-negative (Isl2(-)) RGCs expressing only WT levels of Ephas (see Materials and Methods for numerical values).

**Figure 2.**
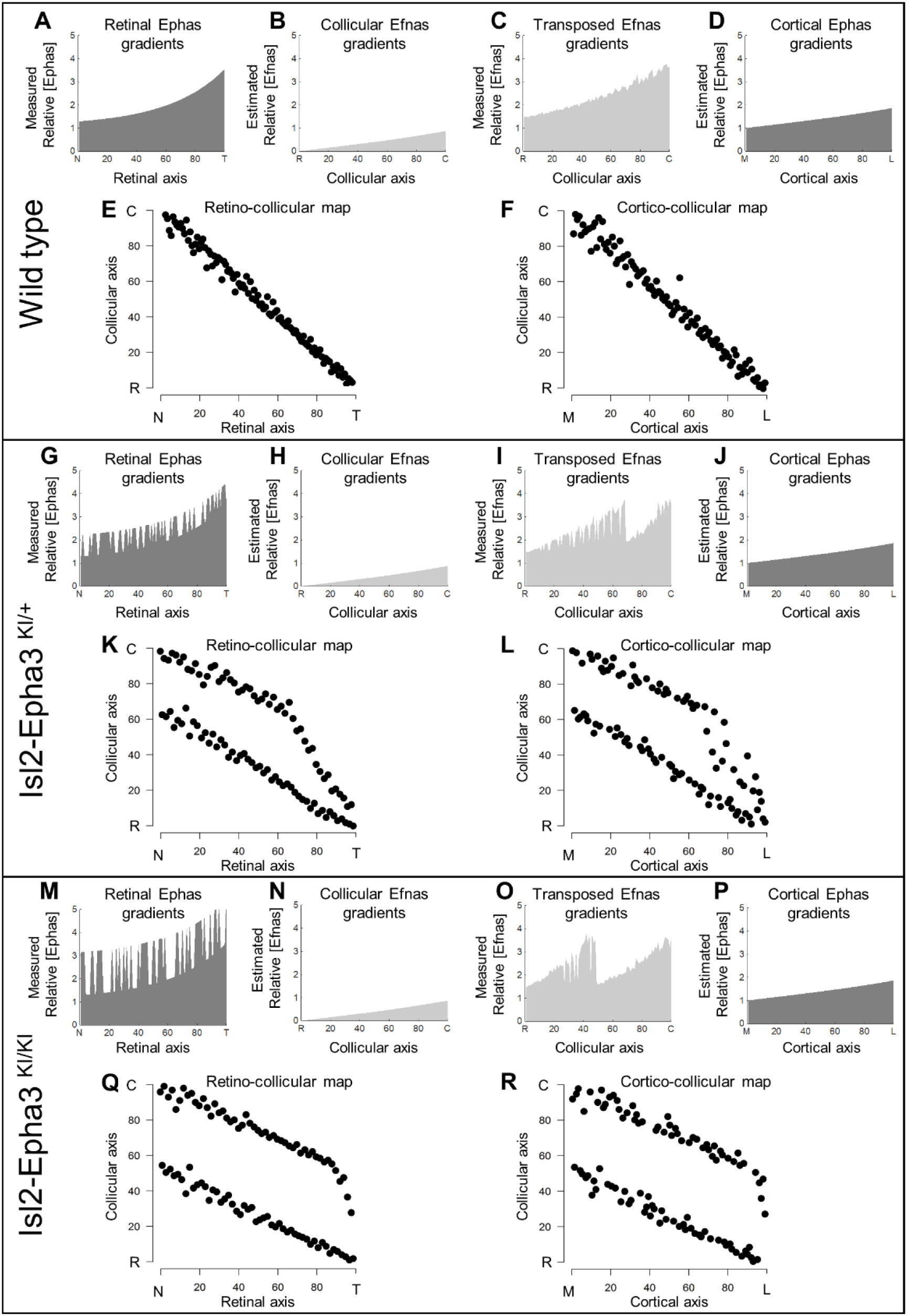
Simulations of retino- and cortico-collicular mapping in Isl2-Epha3KI animals. **A, G, M.** Representation of measured retinal Ephas gradients along the NT axis in WT (A), Isl2-Epha3^KI/+^ (G) and Isl2-Epha3^KI/KI^ (M) animals (see Materials and Methods and Table 1 for equations). **B, H, N.** Representation of the estimated collicular Efnas gradients along the RC axis in WT (B), Isl2-Epha3^KI/+^ (H) and Isl2-Epha3^KI/KI^ (N) animals (see Materials and Methods and Table 1 for equations). **C, I, O.** Representation of the transposed retinal Efnas gradients into the SC along the RC axis in WT (C), Isl2-Epha3^KI/+^ (I) and Isl2-Epha3^KI/KI^ (O) animals (see Materials and Methods and Table 1 for equations). **D, J, P.** Representation of the estimated cortical Ephas gradients along the ML axis in V1 in WT (D), Isl2-Epha3^KI/+^ (J) and Isl2-Epha3^KI/KI^ (P) animals (see Materials and Methods and Table 1 for equations). **E, K, Q.** Simulated retino-collicular map in in WT (E), Isl2-Epha3^KI/+^ (K) and Isl2-Epha3^KI/KI^ (Q) animals generated by the 3-step map alignment algorithm (representative of n = 20 runs). **F, L, R.** Simulated cortico-collicular map in WT (F), Isl2-Epha3^KI/+^ (L) and Isl2-Epha3^KI/KI^ (R) animals generated by the 3-step map alignment algorithm (representative of n = 20 runs). Abbreviations: N, nasal; T, temporal; R, rostral; C, caudal; M, medial; L, lateral.

The projections of one hundred RGCs onto the SC, the retino-collicular map, were simulated by the 3-step map alignment model (10^7^ iterations per run, n = 20 runs) for the WT, Isl2-Epha3^KI/+^ and Isl2-Epha3^KI/KI^ genotypes (Fig 2E, K, Q). RGCs axons/growth cones carrying gradients of Epha receptors are repelled by complementary gradients of collicular Efnas (forward signaling) (Fig 2B, H, N). Results showed a continuous, single map for WT (Fig 2E), as expected from previous experimental findings and theoretical modelling (Brown et al., 2000; Reber et al., 2004; Savier et al., 2017; Bevins et al., 2011). For the Isl2-Epha3^KI/+^, partially duplicated retino-collicular mapping is modelled, together with the presence of a collapse point for temporal-most RGCs (Fig 2K) as extensively observed experimentally (Brown et al., 2000; Reber et al., 2004; Bevins et al., 2011) and theoretically (Reber et al., 2004; Willshaw, 2006; Tsigankov and Koulakov, 2010; Simpson and Goodhill, 2011). For Isl2-Epha3^KI/KI^, the retino-collicular map is fully duplicated, similarly to previous experimental and theoretical findings (Fig 2Q) (Brown et al., 2000; Reber et al., 2004; Willshaw, 2006; Tsigankov and Koulakov, 2010; Bevins et al., 2011; Simpson and Goodhill, 2011). In the following step, the gradient of experimentally measured retinal Efnas is transposed to the SC according to the layout of the retino-collicular map (Fig 2C, I, O). For WT, this generates a smooth monotonically increasing gradient (Fig 2C). For the Isl2-Epha3^KI/+^ genotype, this transposition generates a double oscillatory gradient of retinal Efnas in the SC (Fig 2I), a consequence of the partial duplication of the retino-collicular map (Fig 2K). For the Isl2-Epha3^KI/KI^, the transposition of the retinal Efnas gradient, according to the fully duplicated retino-collicular map (Fig 2Q), generated a double oscillatory gradient of transposed retinal Efnas in the SC (Fig 2O).

Finally, the projections of one hundred V1 neurons onto the SC (the cortico-collicular map) were simulated for all three genotypes (Fig 2F, L, R). V1 axons/growth cones carrying gradients of Epha receptors (Fig 2D, J, P) are repelled by the transposed gradients of retinal Efnas into the SC (forward signaling) as suggested previously (Savier et al., 2017). The WT cortico-collicular map is smooth and continuous (Fig 2F), similarly to the retino-collicular map (Fig 2E). The Isl2-Epha3^KI/+^ simulations show dispersed projections forming two separated maps collapsing in the rostral-most part of the SC from lateral-most V1 neurons (Fig 2L). Cortico-collicular map simulations for the Isl2-Epha3^KI/KI^ genotype shows dispersed projections forming two fully separated maps (Fig 2R).

### Experimental validation of the 3-step map alignment model

To validate the simulations of the model predicting the formation of visual maps in the Isl2-Epha3KI animals, we performed anterograde retino- and cortico-collicular tracings *in vivo*. In the Isl2-Epha3^KI/+^ animals, two experimental measurements (Fig 3A, triangles and square in Fig 3B) are shown, plotted on the simulated retino-collicular map (black dots, grey/black lines, Fig 3B) indicating a duplicated projection for cells from the nasal pole of the retina as extensively shown earlier (Brown et al., 2000; Reber et al., 2004; Savier et al., 2017). For RGCs located on the temporal side of the retina, a single projection can be found as was also described previously (Brown et al., 2000; Reber et al., 2004; Savier et al., 2017). Our experimental measurements (triangles and square, Fig 3B) match with the theoretical representation (black dots, black/grey lines Fig 3B). Further validation of the algorithm was performed by *in vivo* analysis of the cortico-collicular mapping in the Isl2-Epha3^KI/+^ animals. Cortico-collicular anterograde tracing (Fig 3D, n = 15, red dots/lines), performed as described earlier (Savier et al., 2017), revealed a partially duplicated cortico-collicular map with the occurrence of a collapse point near 82% of the medial-lateral axis of V1 (16% of the rostral-caudal axis of the SC), similarly to the simulated map (black dots, black/grey lines Fig 3D, two-samples Kolmogorov-Smirnov test). Two of the injections and their corresponding terminations zones in the SC are depicted (Fig 3C, arrows and arrowhead in Fig 3D).

**Figure 3.**
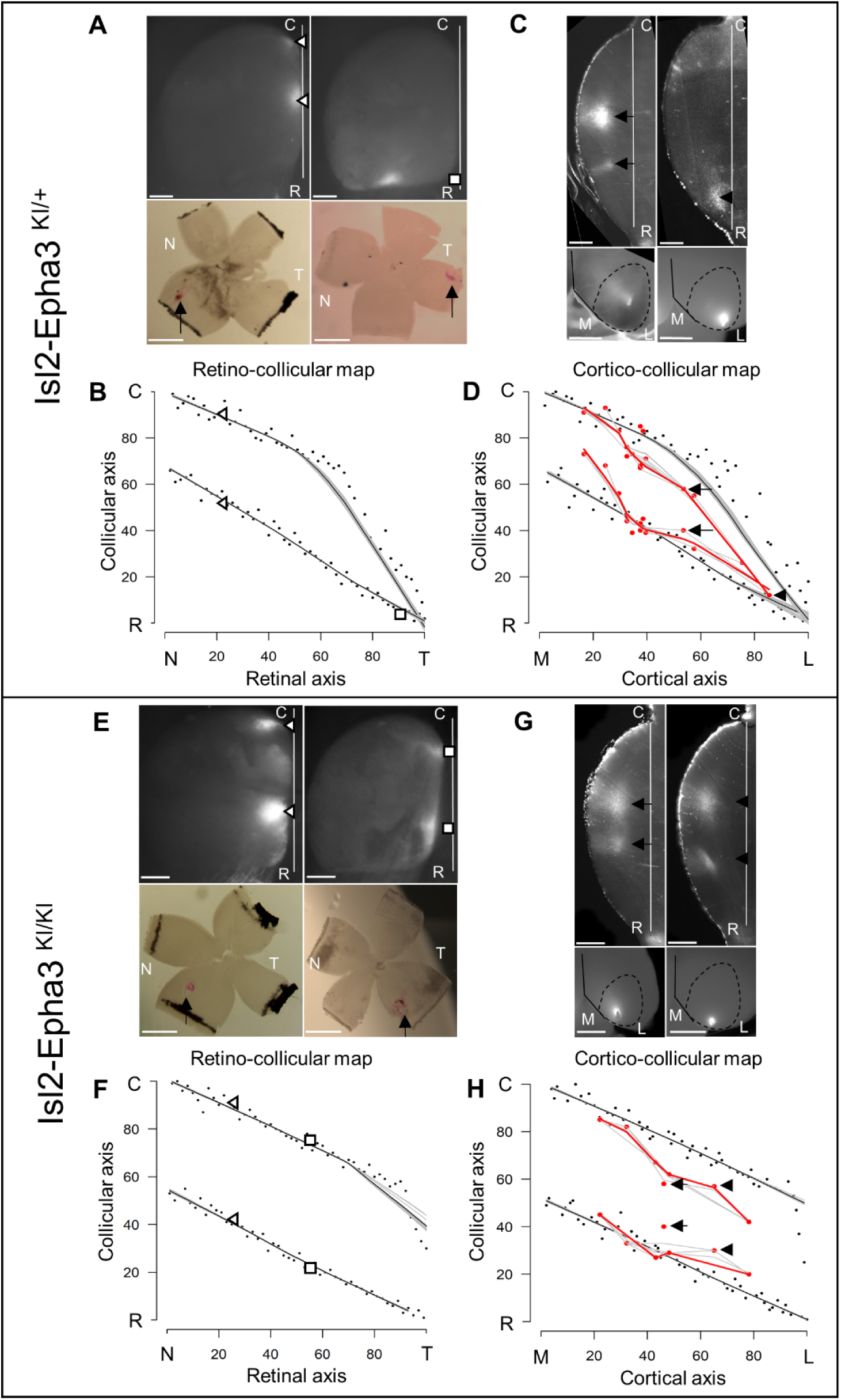
Experimental validation of retino- and cortico-collicular mapping in Isl2-Epha3KI animals. **A.** Images of two experimental injections showing the collicular terminations zones (triangles and square, top-view, upper panels) after focal retinal injections (arrows, flat-mount, lower panel) in Isl2-Epha3^KI/+^ animals. **B.** Cartesian representation of the injections (triangles and square) in (A) superimposed with the simulated retino-collicular map (black dots, n = 100) in Isl2-Epha3^KI/+^. Map profile is calculated by LOESS smoothing (black and grey lines). **C.** Images of two experimental injections showing the collicular termination zones (sagittal view, upper panels) after focal cortical V1 injection (top-view, lower panels). Arrows and arrowheads indicate the site of the termination zones. Lower left panel shows CO staining (dark grey) delineating V1. **D.** Cartesian representation of the experimental (red dots/lines, n = 15 animals) and simulated (black dots, n = 100) cortico-collicular maps calculated by LOESS smoothing (black, red and grey lines). Arrows and arrowhead represent the two injections shown in (C). Two-samples Kolmogorov-Smirnov test, D-stat = 0.273 < D-crit. = 0.282, p = 0.06, simulated and experimentally measured cortico-collicular maps are not significantly different. **E.** Images of two experimental injections showing the collicular terminations zones (triangles and squares, top-view, upper panels) after focal retinal injection (arrows, flat-mount, lower panel) in Isl2-Epha3^KI/KI^ animals. **F.** Cartesian representation of the injections (triangles and squares) in (E) superimposed with the simulated retino-collicular map (black dots, n = 100) in Isl2-Epha3^KI/KI^. Map profile is calculated by LOESS smoothing (black and grey lines). **G.** Images of two experimental injections showing the collicular duplicated termination zones (arrows and arrowheads, sagittal view, upper panels) after focal cortical V1 injection (top-view, lower panels). **H.** Cartesian representation of the experimental (red dots/lines, n = 7 animals) and simulated (black dots, n = 100) cortico-collicular maps calculated by LOESS smoothing (black, red and grey lines). Arrows and arrowheads represent the two examples in (G). Two-samples Kolmogorov-Smirnov test, D-stat = 0.190 < D-crit. = 0.371, p = 0.72, simulated and experimentally measured cortico-collicular maps are not significantly different. Scale bars: 400 μm (A upper, C, E upper, G), 1mm (A, E lower). Abbreviations: N, nasal; T, temporal; R, rostral; C, caudal; M, medial; L, lateral.

In Isl2-Epha3^KI/KI^, retino-collicular duplications are observed as shown by two experimental measurements (triangles and squares) plotted on the retino-collicular map (Fig 3E, F) and as extensively described previously (Brown et al., 2000; Reber et al., 2004). These experimental data match with the simulated mapping (black dots, black/grey lines Fig 3F). Cortico-collicular map tracing *in vivo* (Fig 3G, H, red dots/line, n = 7) revealed a fully duplicated cortico-collicular map, as predicted by the simulations (Fig 3H, black dots, black/grey lines, two-samples Kolmogorov-Smirnov test). Two of the injections and their corresponding terminations zones in the SC are depicted (Fig 3G, arrows and arrowhead in Fig 3H,). These results further validate the predictions and reinforce the underlying mapping principle encoded in the 3-step map alignment algorithm.

### Retino/cortico-collicular maps organization indexes

Previous work demonstrated that retino- and cortico-collicular maps must be aligned and in register to allow efficient detection of visual stimuli by the SC (Zhao et al., 2014; Liang et al., 2015; Basso and May, 2017). We further implemented the 3-step map alignment algorithm to calculate an “intrinsic dispersion index” (IDI) for each visual map (IDI_retino_ and IDI_cortico_) and a “alignment index” (AI) for each genotype. These indexes measure the overall organization of the retino- and cortico-collicular maps. IDI_retino_ and IDI_cortico_ indicate the degree of dispersion of the corresponding map (or within-map variability) and should be minimum. AI represents the degree of alignment between the retino- and cortico-collicular maps and should be close to 1. AI = 1 corresponds to the theoretical one-to-one alignment of all the RGCs projections with all the V1 projections onto the SC, however such value will not be observed due to the intrinsic variability of the mapping introduced by the stochastic process of spontaneous activity. In WT animals, median IDI_retino_ = 5.66, 95% CI [5.15; 6.17], median IDI_cortico_ = 8.12 [7.63; 8.60] and median AI = 2.23 [2.13; 2.33] (Fig 4A, G). These values correspond to the control values for aligned, single retino- and cortico-collicular maps (Fig 4B, black and white dots). In Isl2-Epha3^KI/+^, median IDI_retino_ = 38.8 [36.8; 40.8], median IDI_cortico_ = 38.0 [36.1; 40.0] and median AI = 2.07 [1.95; 2.19] (Fig 4A) indicating increased spreading of both retino- and cortico-collicular projections, due to the partially duplicated maps (Fig 4C, black and white dots). The median AI value in Isl2-Epha3^KI/+^ (2.07 [1.95, 2.19]) is not statistically different from WT (Fig 4G), suggesting that Isl2-Epha3^KI/+^ retino/cortico-collicular maps are aligned (Fig 4C, black and white dots). In the Isl2-Epha3^KI/KI^ animals, median IDI_retino_ = 51.7 [50.8; 52.6], median IDI_cortico_ = 50.6 [49.7; 51.5] and median AI = 2.2 [2.08; 2.32] (Fig 4A, G), indicating an increased separation between the retino- and cortico-collicular maps compared to WT and Isl2-Epha3^KI/+^ due to fully duplicated projections (Fig 4D white and black dots). The median AI value (2.20 [2.02; 2.43]) is not significantly different from WT (Fig 4G), indicating aligned retino- and cortico-collicular maps (Fig 4D, black and white dots). Median AI values for WT, Isl2-Epha3^KI/+^ and Isl2-Epha3^KI/KI^ are not significantly different suggesting that retino- and cortico-collicular maps in these animals are aligned (Fig 4G), although partially or fully duplicated.

**Figure 4.**
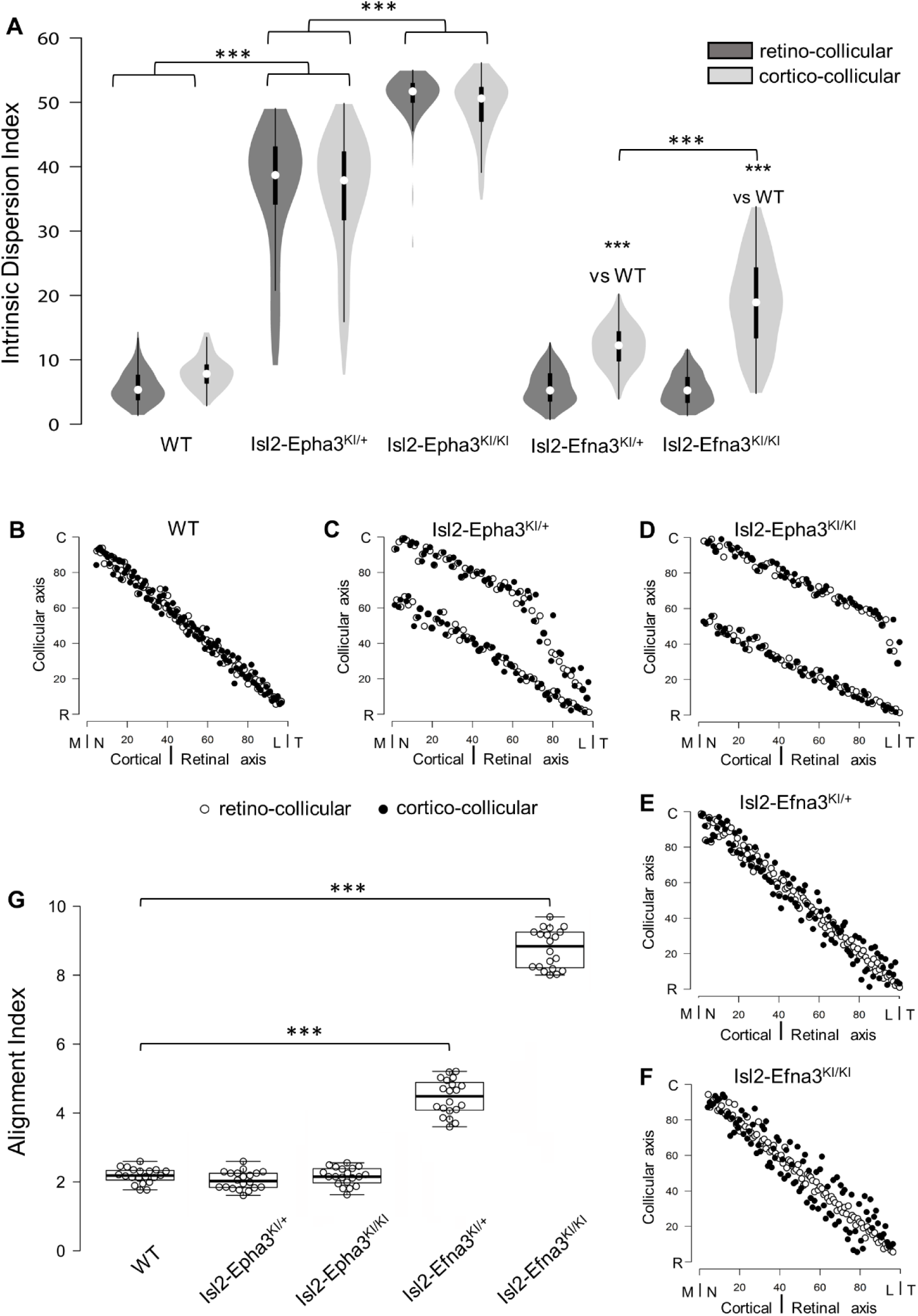
Intrinsic dispersion index (IDI) and alignment index (AI) in Isl2-Epha3KI and Isl2-Efna3KI animal models. **A.** Violin plot representation of the median IDIs (from n = 10 simulated maps, each composed of 100 projections) in WT, Isl2-Epha3KI and Isl2-Efna3KI animal models. Mann-Whitney test: WT IDI_retino_ vs. Isl2-Epha3^KI/+^ IDI_retino_, z-score = 12.08, effect r = 0.85, p = 6E-55; WT IDI_cortico_ vs. Isl2-Epha3^KI/+^ IDI_cortico_, z-score = 12.04, effect r = 0.85, p = 8E-52; Isl2-Epha3^KI/+^ IDI_retino_ vs Isl2-Epha3^KI/KI^ IDI_retino_, z-score = 11.25, effect r = 0.80, p = 2E-39; Isl2-Epha3^KI/+^ IDI_cortico_ vs Isl2-Epha3^KI/KI^ IDI_cortico_, z-score = 10.53, effect r = 0.74, p = 1E-32; WT IDI_cortico_ vs Isl2-Efna3^KI/+^ IDI_cortico_, z-score = 8.30, effect r = 0.59, p = 1E-18; WT IDI_cortico_ vs Isl2-Efna3^KI/KI^ IDI_cortico_, z-score = 10.37, effect r = 0.73, p = 3E-31; Isl2-Efna3^KI/+^ IDI_cortico_ vs Isl2-Efna3^KI/KI^ IDI_cortico_, z-score = 6.93, effect r = 0.49, p = 6E-13; *** p < 0.001. **B, C, D, E, F.** Representation and superimposition of simulated retino-collicular (white dots) and cortico-collicular (black dots) maps in WT (B), Isl2-Epha3^KI/+^ (C), Isl2-Epha3^KI/KI^ (D), Isl2-Efna3^KI/+^ (E) and Isl2-Efna3^KI/KI^ (F) animals (representative of n = 10 runs). **G.** Box plot representation of median AI (from n = 20 simulated retino/cortico-collicular maps) in WT, Isl2-Epha3^KI/+^, Isl2-Epha3^KI/KI^, Isl2-Efna3^KI/+^ and Isl2-Efna3^KI/KI^ animals. Mann-Whitney test: AI WT vs. AI Isl2-Epha3^KI/+^, z-score = 1.62, effect r = 0.26 p = 0.10; AI WT vs. AI Isl2-Epha3^KI/KI^, z-score = 0.11, effect r = 0.02, p = 0.90; AI WT vs AI Isl2-Efna3^KI/+^, z-score = 5.40, effect r = 0.85, p = 1.45E-11; AI WT vs AI Isl2-Efna3^KI/KI^, z-score = 5.40, effect r = 0.85, p = 1.45E-11. *** p < 0.001. Abbreviations: IDI, intrinsic dispersion index; AI, alignment index; WT, wildtype; N, nasal; T, temporal; R, rostral; C, caudal; M, medial; L, lateral.

To further test and validate these map organization indicators, we calculated the IDIs and AI in previously characterized Isl2-Efna3^KI/+^ and Isl2-Efna3^KI/KI^ animals (Savier et al., 2017). Median values for Isl2-Efna3^KI/+^ are IDI_retino_ = 5.56 [5.04; 6.07] and IDI_cortico_ = 12.5 [11.8; 13.2] (Fig 4A). IDI_retino_ is not significantly different from WT suggesting no dispersion of the retino-collicular map (Fig 4E, white dots). However, IDI_cortico_ is significantly different from WT indicating a spreading of the cortico-collicular map (Fig 4E, black dots). This suggests a mis-alignment between retino- and cortico-collicular maps (Fig 4E) as confirmed by the median AI value in Isl2-Efna3^KI/+^ (Fig 4G, 4.52 [4.28, 4.76]), significantly different from WT. Visual maps mis-alignment is more pronounced in Isl2-Efna3^KI/KI^ (median AI = 8.85 [8.23, 9.26], Fig 4G) compared to Isl2-Efna3^KI/+^ as previously shown *in vivo* (Savier et al., 2017). Map dispersion value IDI_retino_ in Isl2-Efna3^KI/KI^ indicates that retino-collicular map spreading is similar to WT (Fig 4F, white dots; Fig 4A, median IDI_retino_ = 5.56 [5.08, 6.03], not significantly different from WT) whereas IDI_cortico_ indicates dispersion of the cortico-collicular map (Fig 4F, black dots; Fig 4A, median IDI_cortico_ = 19.1 [17.7, 20.5]) as demonstrated previously (Savier et al., 2017). Such significant dispersion of the cortico-collicular map leads to an important misalignment between retino- and cortico-collicular projections in Isl2-Efna3^KI/KI^ animals (median AI = 8.85 [8.59; 9.11] significantly different from WT, Fig 4F, G).

We undertook a more detailed analysis of the mapping organization and alignment by implementing the local “intrinsic dispersion variation” (IDV) (y axis) for each retino- and cortico-collicular maps along the rostral-caudal axis of the SC (x axis, Fig 5). Median local IDVs indicate the degree of dispersion and alignment as a function of the position along the rostral-caudal axis in the SC. In WT (Fig 5A), Isl2-Epha3^KI/+^ (Fig 5B) and Isl2-Epha3^KI/KI^ (Fig 5C), median local IDVs between retino- and cortico-collicular maps are similar and covary (> 0) (WT retino/cortico: Jaccard similarity index = 0.60 -indicating 60% overlap-covariance = 1.64; Isl2-Epha3^KI/+^ retino/cortico: Jaccard similarity index = 0.30, covariance = 88.2; Isl2-Epha3^KI/KI^ retino/cortico: Jaccard similarity index = 0.55, covariance = 12.8), indicating that the maps are aligned. This is in sharp contrast to the Isl2-Efna3^KI/+^ (Fig 5D) and Isl2-Efna3^KI/KI^ (Fig 5E) animals where median local retino- and cortico-collicular IDVs variation do not superimpose nor covary (Isl2-Efna3^KI/+^ retino/cortico: Jaccard similarity index = 0.17, covariance = -1.99; Isl2-Efna3^KI/KI^ retino/cortico: Jaccard similarity index = 0.08, covariance = -4.13). Moreover, values of IDV are indicative of map dispersion along the RC axis in the SC. We calculated a cut-off value of median local IDV based on WT retino/cortico-collicular maps organization (median local IDV_threshold_ = 14.7, the minimum local IDV, dashed black line in Fig 5) corresponding to the WT maps dispersion. Values greater than the local IDV_threshold_, indicate a duplication of the map, while values below this threshold indicate a single map. In WT (Fig 5A, yellow), Isl2-Efna3^KI/+^ (Fig 5D, light red) and Isl2-Efna3^KI/KI^ (Fig 5E, light blue), median local IDV_retino_ representations suggest a similar dispersion along the RC axis with all IDV values below IDV_threshold_ indicating single maps and corresponding to experimental observations (Savier et al., 2017). In Isl2-Epha3^KI/+^ IDV_retino_ and IDV_cortico_ (Fig 5B), Isl2-Epha3^KI/KI^ IDV_retino_ and IDV_cortico_ (Fig 5C), Isl2-Efna3^KI/+^ IDV_cortico_ (Fig 5D, dark red) and Isl2-Efna3^KI/KI^ IDV_cortico_ (Fig 5E, dark blue), values are above IDV_threshold_, indicating map duplication. More precisely, in Isl2-Epha3^KI/+^ animals, median local IDV_retino_ and median local IDV_cortico_ reach IDV_threshold_ at 12% and at 8% of the rostral-caudal axis in the SC respectively (Fig 5B) consistent with the position of the collapse points described earlier (Brown et al., 2000; Savier et al., 2017 and Fig 3B, D). This suggests single retino- and cortico-collicular maps in the rostral-most pole of the SC and a duplicated visual maps from 8%-12% to 100% of the rostral-caudal axis as shown experimentally and theoretically (Brown et al., 2000; Savier et al., 2017; Fig 3B, D). In Isl2-Epha3^KI/KI^ animals, both retino- and cortico-median local IDV > IDV_threshold_ along the entire rostral-caudal axis (Fig 5C), indicating full retino- and cortico-collicular maps duplication as shown experimentally (Brown et al., 2000; Triplett et al., 2009; Fig 3F, H). Finally, for both Isl2-Efna3^KI/+^ and Isl2-Efna3^KI/KI^ cortico-collicular maps (Fig 5D, dark red, Fig 5E, dark blue, respectively) local IDVs oscillate around the IDV_threshold_ with a more pronounced effect in the rostral-half of the SC in Isl2-Efna3^KI/KI^ indicating partial duplications of the cortico-collicular maps in these animals, as demonstrated *in vivo* (Savier et al., 2017). Retino-collicular median local IDVs in Isl2-Efna3^KI/+^ and Isl2-Efna3^KI/KI^ animals (Fig 5D pink line, Fig 5E light blue line, respectively) fall below IDV_threshold_ indicating single maps as demonstrated previously (Savier et al., 2017). Altogether these results suggest that IDIs, AIs and median local IDVs are relevant, reliable and accurate indicators of visual maps organization and conformation.

**Figure 5.**
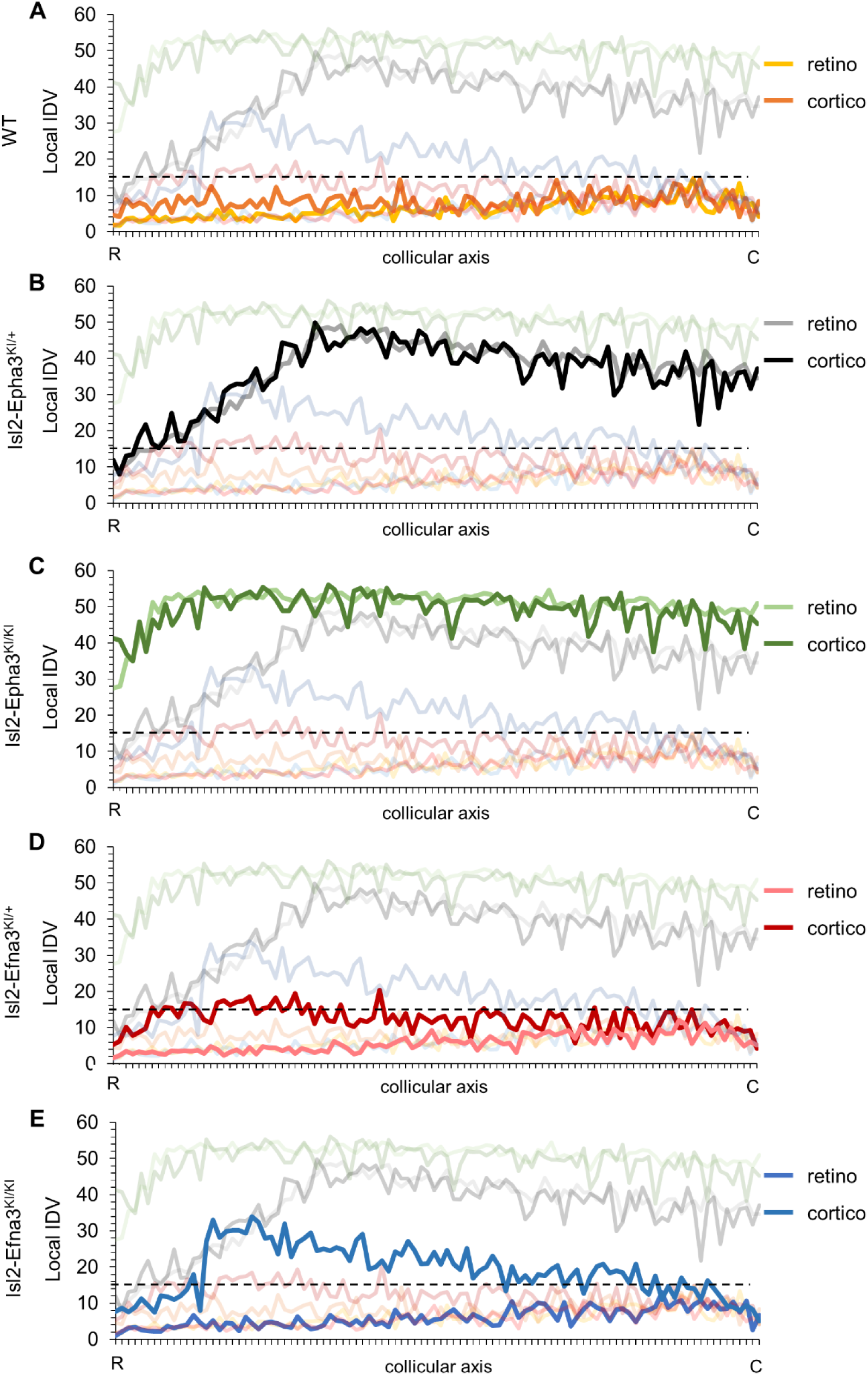
Local intrinsic dispersion variation (local IDV) in WT, Isl2-Epha3KI and Isl2-Efna3KI animal models. **A, B, C, D, E.** Representation of the local IDV values for both retinal (light colors) and cortical (dark colors) projections along the rostral R (0%) – caudal C (100%) axis of the SC in WT (A), Isl2-Epha3^KI/+^ (B), Isl2-Epha3^KI/KI^ (C), Isl2-Efna3^KI/+^ (D) and Isl2-Efna3^KI/KI^ (E) animals (representative of n = 10 runs). Dashed line represents the threshold above which maps are duplicated. Abbreviations: IDV, intrinsic dispersion variation; WT, wildtype; R, rostral; C, caudal.

## Discussion

### The 3-step map alignment model replicates other visual map-defective mutants

Taken together, these experimental and theoretical results confirm the validity and robustness of the 3-step map alignment algorithm in predicting retino- and cortico-collicular maps formation and alignment during development. These findings broaden the simulation capacity of the model to the formation and alignment of visual maps in Isl2-Epha3KI mutant animals originally described in 2000 (Brown et al., 2000). After 10^7^ iterations per run, the algorithm generates retino-collicular maps for Isl2-Epha3^KI/+^ heterozygous and Isl2-Epha3^KI/KI^ homozygous animals. In Isl2-Epha3^KI/KI^, the simulated retino-collicular map is fully duplicated, as described *in vivo* (Brown et al., 2000; Reber et al., 2004; Fig 2Q, 3F) whereas in the Isl2-Epha3^KI/+^, the simulated retino-collicular duplication collapses at approximately 80% of the nasal-temporal axis, similarly to previous results (Brown et al., 2000; Reber et al., 2004; Fig 2K, 3B). More recent experimental analyses show a collapse point appearing within a range of 74% to 80% of the nasal-temporal axis (Savier et al., 2017) in contrast to previous results indicating a collapse at 76% of the nasal-temporal axis (Brown et al., 2000; Reber et al., 2004). This discrepancy might be explained by the measurement methods which differ between the original Brown et al. publication (Brown et al., 2000) and the recent Savier et al. (Savier et al., 2017) and here (Fig 3B, D, E, F). In the last two, the Locally Weighted Scatterplot Smoothing (LOWESS) cross validation method (Efron and Tibshirani, 1991) was used on both experimental and simulated maps providing a range of values, instead of a given value as performed in Brown et al. (Brown et al., 2000), for the occurrence of the collapse point along the nasal-temporal axis (see also Fig 6 Sup. Fig 1 in Savier et al., 2017). In Isl2-Epha3KI animals, the lower retino-collicular maps corresponds to Isl2(+) Epha3-expressing RGCs, covering 0% to 50% of the rostral-caudal axis in the Isl2-Epha3^KI/KI^ and 0% to 80% of the axis in Isl2-Epha3^KI/+^ (Fig 5, 6 in Brown et al., 2000). The upper map, corresponding to the Isl2(-) WT RGCs, covers the caudal half (50% to 100%) of the rostral-caudal axis of the SC in Isl2-Epha3^KI/KI^ and 20%-100% of the axis in Isl2-Epha3^KI/+^ (Fig 5, 6 in Brown et al., 2000). In both contexts, nasal Isl2(-) RGCs, expressing high levels of Efnas, project to caudal locations in the SC whereas nasal Isl2(+) RGCs, also carrying high levels of Efnas, project ectopically in a more rostral part of the SC, where the WT levels of retinal Efnas are normally low (compare Fig 2C with Fig 2I and 2O; see also Fig 6). Such distribution of RGC projections in the SC in both Isl2-Epha3^KI/KI^ and Isl2-Epha3^KI/+^ animals leads to a perturbation of the transposed retinal Efnas gradient in the SC, generating duplicated Efnas gradient along the rostral-caudal axis in both genotypes (Fig 2I, O, Fig 6). Consequently, simulated cortico-collicular maps are also duplicated for both Isl2-Epha3^KI/+^ and Isl2-Epha3^KI/KI^ and further align with the duplicated retino-collicular maps as confirmed by *in vivo* analyses (Fig 3D, H, Fig 6).

**Figure 6.**
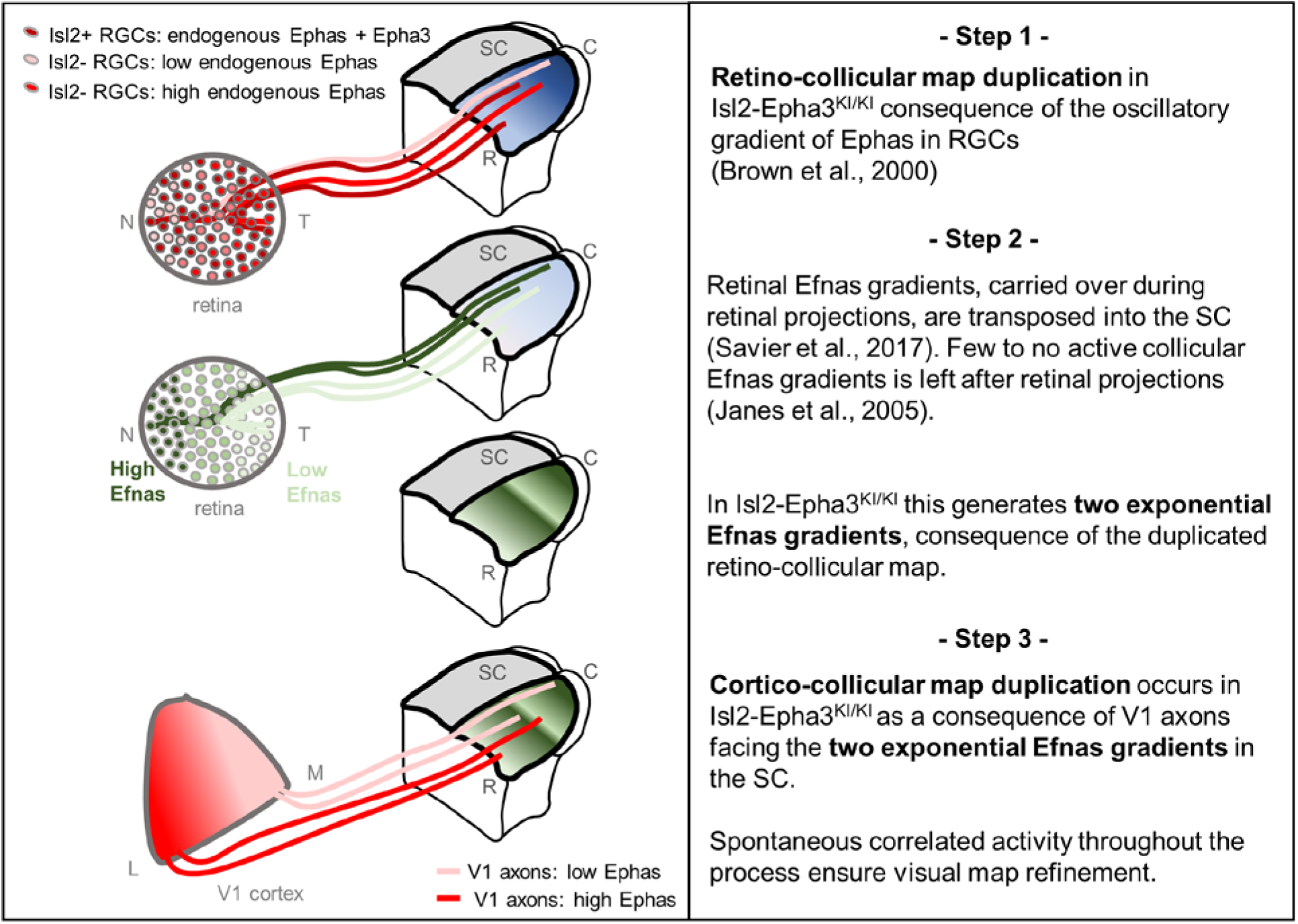
Proposed mechanism of visual maps duplication and alignment in Isl2-Epha3KI animals based on the 3-step map alignment model. **Step 1:** Isl2(+) RGCs expressing WT levels Ephas + ectopic Epha3 (dark red) and Isl2(-) RGCs expressing WT levels of Ephas (light red to red) send their axons into the SC during the first postnatal week. These retino-collicular projections form a duplicated map due to the oscillatory gradient of Ephas receptors in the RGCs reading the WT collicular Efnas gradients (blue R-C gradient in SC) through forward signaling. **Step 2:** Retinal Efnas gradients (high Efnas-nasal-dark green, low Efnas-temporal-light green) are carried to the SC during the formation of the retino-collicular projections. This transposition of retinal Efnas generates two exponential gradients of Efnas in the SC, due to the duplication of the retino-collicular map and replaces the WT collicular Efnas gradients previously used by the RGCs axons (Janes et al., 2005) **Step 3:** V1 axons, expressing smooth gradients of Ephas receptors (light red – red), are facing two exponential gradients of Efnas, of retinal origin, in SC. Through forward signaling, this 2 exponential Efnas gradients generate a duplication of the cortico-collicular projections, which aligns with the retino-collicular map. Abbreviations: N, nasal; T, temporal; M, medial; L, lateral; R, rostral; C, caudal; SC, superior colliculus; RGCs, retinal ganglion cells

### The basic maps organization principle encoded in the 3-step map alignment model provides an alternative explanation for map alignment defects in Isl2-Epha3KI animals

Our quantification of Epha and Efna throughout the visual system revealed that the ectopic over-expression of Epha3 in the retina has no effect on the expression of Efnas in this structure. These findings exclude a potential alteration of the endogenous gradient of Efnas in the retina. Previous studies (Triplett et al., 2009) suggested that the redistribution of the correlated activity was responsible for the duplication of the cortico-collicular map. Our results suggest an alternative, although complementary mechanism, in which the redistribution of the molecular cues carried by retinal axons during the formation of a duplicated retino-collicular map is sufficient to induce a collicular duplication of the projections coming from V1 (Fig. 6). As mentioned earlier (Savier et al., 2017), in this context retinal Efnas, free or bound to Ephas to a limited amount (Suetterlin and Drescher, 2014), retain their binding activity in the SC for incoming V1 axons carrying Epha receptors. Since the retino-collicular map is duplicated in the Isl2-Epha3KI, the nasal-temporal axis of the retina, along which the retinal Efna gradient runs, is represented twice in the collicular space. This rearrangement of the retinal Efnas transposed to the SC generates a duplicated gradient which is then encountered by the incoming cortical V1 axons. At this point in development, the endogenous collicular Efnas are not available anymore as, for the most part, they have been endocytosed upon trans-binding with Epha receptors on retinal axons (Janes et al., 2005; Savier et al., 2017). As a consequence, the optimal local amount of transposed Efnas signalling the corresponding retinotopic position to V1 axons exists at two distinct locations along the rostral-caudal axis, leading to duplication of the cortico-collicular map. Our previous findings in the characterization of the Isl2-Efna3KI mutant revealed an incomplete penetrance of the cortico-collicular duplication phenotype, suggesting a counter-balancing role of the correlated activity within the normal retino-collicular map. For the Isl2-Epha3KI, the correlated activity is also duplicated along the rostro-caudal axis, due to the duplicated retino-collicular map, reinforcing this “duplication” effect and leading to a stronger penetrance of the phenotype. To further test this hypothesis, it would be interesting to selectively manipulate correlated activity during development in either the retina, the SC or V1. Our current conceptual framework suggests that altering the activity in the SC would still result in a duplication of the cortico-collicular map of the Isl2-Epha3KI and an increase in the amount of cortico-collicular duplication in the Isl2-Efna3KI. Moreover, mapping and alignment along the lateral-medial axis of the SC involving EphB/Efnb signalling is not addressed here as the Isl2-Epha3KI animals only present mapping defect along the rostral-caudal axis (Brown et al., 2000; Triplett et al., 2009).

### Quantifying the degree of maps alignment and organization

We performed a quantitative analysis of the degree of visual maps organization, implementing the intrinsic dispersion index (IDI) for each visual map, the mean alignment index (AI) (Fig 4) and the local intrinsic dispersion variation (local IDV) (Fig 5). These indexes objectively describe and quantify the degree of organization and conformation of both retino- and cortico-collicular maps, providing a detailed measure of the dispersion of each map and a measure of the degree of alignment between the maps as a whole and locally. In general, the experimental number of measured projections is low, approximately n = 15 to 20 total, compared to the n = 100 (or more) projections generated by the algorithm. This suggests that, if the algorithm accurately simulates maps organization, the indicators, IDI, AI and local IDVs, inferred from this algorithm are more precise and refined than those inferred from experimental maps. As demonstrated above, the 3-step map alignment algorithm simulates and predicts visual map defects in Isl2-Epha3KI animals as well as in other animal models (Savier et al., 2017) indicating its reliability and robustness; therefore, we inferred the IDIs AIs and local IDVs from the algorithm-generated maps for each genotype. An increase of both IDIs and local IDVs in Isl2-Epha3^KI/+^ and Isl2-Epha3^KI/KI^ animals correlates with increased spreading of the visual maps, consequence of the duplicated retino- and cortico-collicular projections. By contrast, AI values in these animals are similar to WT, indicating alignment of the retino- and cortico-collicular maps, as observed *in vivo*. The relevance of the 3-step map alignment algorithm and the validity of the map organization indexes are further confirmed by the analysis of map organization in the previously characterized Isl2-Efna3KI animals. Here, IDIs, AIs and local IDVs generated by the algorithm predict visual map organization as previously shown *in vivo* (Savier et al., 2017). High AI values (> 2.5 = WT median + 95% CI) indicate a significant misalignment between retino- and cortico-collicular projections, in particular in Isl2-Efna3^KI/KI^ (confirmed by the low Jaccard similarity index and negative covariance). Such misalignment is also indicated by a significant difference in dispersion (IDI) of the cortico-collicular map when compared to the retino-collicular map (IDI_retino_ ≠ IDI_cortico_). These observations are further validated and confirmed by the local IDVs graph (Fig 5) which delivers detailed information about the conformation of the maps along the rostral-caudal axis of the SC. Descriptive analyses of the IDVs using Jaccard similarity index and covariance guaranties robust data as to the degree of similarity of the maps (Metcalf and Casey, 2016). The implementation of these indexes provides a very detailed and robust analysis of the layout and a qualitative measure of the organization and alignment of the visual maps in different mice genetic models. It would also be interesting to characterize other compound mutants which present more separated maps but high alignment like the Isl2-Epha3KI/ Epha4KO (Reber et al., 2004) or the Isl2-Epha3KI/ Epha5KO (Bevins et al., 2011) double mutants.

## Materials and Methods

### Retinal ganglion cell isolation

P1/P2 retinas were freshly dissected and RGCs were isolated and purified (>99%). For details see (Steinmetz et al., 2006) and (Claudepierre et al., 2008). Briefly, cells were harvested in Neurobasal medium (Gibco/Invitrogen) supplemented with (all from Sigma, except where indicated) pyruvate (1 mM), glutamine (2 mM; Gibco/Invitrogen), N-acetyl-l-cysteine (60 mg/ml), putrescine (16 mg/ml), selenite (40 ng/ml), bovine serum albumin (100 mg/ml; fraction V, crystalline grade), streptomycin (100 mg/ml), penicillin (100 U/ml), triiodothyronine (40 ng/ml), holotransferrin (100 mg/ml), insulin (5 mg/ml) and progesterone (62 ng/ml), B27 (1:50, Gibco/Invitrogen), brain-derived neurotrophic factor (BDNF; 25 ng/ml; PeproTech, London, UK), ciliary neurotrophic factor (CNTF; 10 ng/ml; PeproTech) and forskolin (10 mM; Sigma). After isolation, RGCs were treated for RNA extraction.

### Quantitative RT-PCR

V1 cortices, superficial layers of the SC and retinas were freshly dissected. Retinas were cut in three equal pieces along the NT axis (Nasal, Central, Temporal RGCs) and RGCs acutely isolated (Steinmetz et al., 2006; Claudepierre et al., 2008; see above). Total RNA was extracted and quantified as previously described (Savier et al., 2017). Briefly, relative quantification was performed using the comparative Delta Ct method. Duplicates were run for each sample and concentrations for the target gene and for two housekeeping genes (hypoxanthine-guanine phosphoribosyl transferase - Hprt and glyceraldehyde 3-phosphate dehydrogenase – Gapdh) were computed.

### Projections analysis/mapping

Anterograde DiI (1,1-dioctadecyl-3,3,3,3-tetramethylindocarbocyanine perchlorate) or DiD (1,1’– dioctadecyl-3,3,3’,3’-tetramethylindodicarbocyanine, 4-chlorobenzenesulfonate) labelling were performed blind to genotype as described (Savier et al., 2017). Retinas were dissected and imaged using Leica macroscope (MG0295) and LASAF software. Retino-collicular projection coordinates of the DiI injections were calculated as described (Brown et al., 2000; Reber et al., 2004; Bevins et al., 2011). For cortico-collicular map analyses, sagittal vibratome sections were performed on P14 SC and terminal zones (TZs) were plotted along the rostral-caudal axis on Cartesian coordinates (y axis). V1 were photographed as whole-mount and focal injections plotted along the V1 lateral-medial axis (x axis) (Savier et al., 2017; Triplett et al., 2009). Cortico-collicular maps were generated using non-parametric smoothing technique, termed LOESS smoothing (Efron and Tibshirani, 1991), to estimate the profile of the one-dimensional mapping from V1 to SC. To estimate the variability in a mapping containing N data points, we repeated the procedure N times with N-1 datapoints, each time dropping a different datapoint. This is termed a ‘leave-one-out’ method and was used in the R Project for Statistical Computing (RRID:SCR_001905). V1 area was determined by CO staining as described (Savier et al., 2017; Zembrzycki et al., 2015) and shown on Figure 3C, lower left panel. All animal procedures were in accordance with national (council directive 87/848, October 1987), European community (2010/63/EU) guidelines and University Health Network. Official agreement number for animal experimentation is A67-395, protocol number 01831.01 and AUP 6066.4 (M.R).

### In silico simulations of the retino- and cortico-collicular maps

In the 3-step map alignment model, Ephas/Efnas forward signaling level is modelled by experimentally measured and estimated values of graded expression levels of Epha receptors (Epha3/4/5/6/7) and Efna ligands (Efna2/3/5) in RGCs, SC and V1 cortex, as previously described (Tsigankov and Koulakov, 2006; Savier et al., 2017). Competition, where two RGCs/V1 neurons cannot project to the same target in the SC, is modelled by indexation of 100 RGCs/V1 neurons projecting to 100 positions in the SC (Koulakov and Tsigankov, 2004; Tsigankov and Koulakov, 2006; Savier et al., 2017). Contribution of correlated neuronal activity -assuming Hebbian plasticity between RGC/V1 axons and collicular neurons in the SC- is modelled by pair-wise attraction inversely proportional to the distance between two RGC/V1 neurons (Savier et al., 2017). The 3-step map alignment algorithm first simulates the projection of 100 RGCs along the nasal-temporal axis of the retina mapping onto the rostral-caudal axis of the SC (the retino-collicular map).

Second, according to the layout of this map, retinal Efna gradients are transposed along the rostral-caudal axis of the SC. Third, the projections of 100 V1 neurons following the medial-lateral axis of V1 cortex are simulated onto the rostral-caudal axis of the SC. Forward Ephas/Efnas signaling is applied for both retino-collicular and cortico-collicular projections.

The 3-step map alignment model in MATLAB (RRID:SCR_001622) (Savier et al., 2017) was used to simulate the formation of both the retino- and cortico-collicular maps in the presence of an oscillatory Epha gradient in the retina. Each brain structure (retina, SC, V1) is modelled as a 1-d array of 100 neurons (N) in each network. Two maps are generated: first, the map from retina to SC; second, the map from V1 to SC. Each map is modelled sequentially in the same way. This model consists in the minimization of affinity potential (E) which is computed as follows:

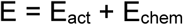

At each step, this potential is minimized by switching two randomly chosen axons probabilistically to reduces the energy in the system by Delta E (ΔE). The probability of switching, p, is given by:

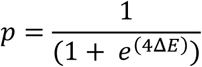

E_chem_ is expressed as follows:

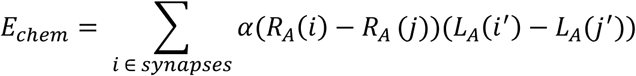

where α= 200 is the strength, R_A_(i) and R_A_(j) the Epha receptor concentration in the retina and V1 at location (i) and (j) and L_A_(i’) and L_A_(j’) the Efna ligand concentration at the corresponding position (i’) and (j’) in the SC.

The contribution of activity-dependent process is modelled as:

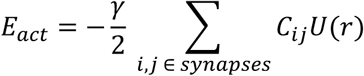

where γ = 1 is the strength parameter, C_ij_ is the cross-correlation of neuronal activity between two RGCs (or V1) neurons during spontaneous activity located in (i) and (j), and U simulates the overlap between two SC cells. Here, we use 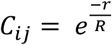, where r is the retinal distance between RGC (i) and (j), R = b x N, with b=0.11 and 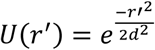, where r’ is the distance between two SC points (i’, j’), d = 3 and N = 1 to 100 neurons. Parameters of the model are presented Table 1.

**Table 1.** summary of the parameters of the 3-step map alignment algorithm.

| Receptor | Epha3 | Epha4 | Epha5 | Epha6 | Source |
| --- | --- | --- | --- | --- | --- |
| Retina | WT = 0<br>Epha3<sup>KI/KI</sup> = 1.86<br>Epha3<sup>KI/+</sup> = 0.93 | 1.05 | 0.14e<sup>0.018x</sup> | 0.09e<sup>0.029x</sup> | Measured (Brown et al., 2000; Reber et al., 2004) |
| V1 | $e^{(-x/100)} - e^{((x - 200)/100) + 1}$ | | | | Estimated (Tsigankov and Koulakov, 2010, 2006) |
| Ligand | EfnA2 | EfnA3 | EfnA5 |  |  |
| Retina | 1.85 e<sup>-0.008x</sup> | 0.44 | 1.79 e<sup>-0.014x</sup> |  | Measured (Savier et al., 2017) |
| SC | $e^{((x - 100)/100)} - e^{((-x-100)/100)}$ | | | | Estimated (Cang et al., 2005; Savier et al., 2017; Tsigankov and Koulakov, 2010, 2006) |
| Parameters |  |  |  |  |  |
| $\gamma$ | 1 | | Strength of activity interaction | | |
| $\alpha$ | 200 | | Chemical strength | | |
| d | 3 |  | SC interaction distance |  |  |
| b | 0.11 |  | Retinal correlation distance |  |  |

### Gradients of ligands and receptors

#### Retinal Epha gradients

Measured gradients of Epha receptors (R _Epha_) in RGCs along the nasal-temporal axis (x) R _Epha_ (x)^retina^ (Figure 2A, G, M, Brown et al., 2000; Reber et al., 2004) are modelled by two equations, one corresponding to Isl2(+) RGCs expressing WT levels of Ephas + Epha3 and the second corresponding to Isl2-negative (Isl2-) RGCs expressing the WT level of Ephas only:

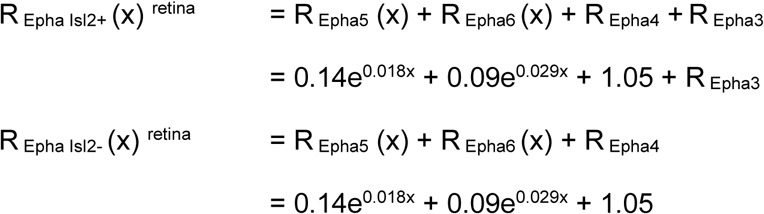

With R _Epha3_ = 0.93 in Isl2-Epha3^KI/+^ and R _Epha3_ = 1.86 in Isl2-Epha3^KI/KI^, since Epha3 expression level depend on the number of copies of the knocked-in allele (Brown et al., 2000; Reber et al., 2004). The oscillatory gradient was generated by randomly attributing to 50% of collicular TZs an over-expression of Epha3 in a genotype-dependent manner.

#### Collicular Efnas gradient

Estimated gradients of Efna ligands (L_Efna_) in the SC along the rostral-caudal axis (x), L_Efna_ (x)^SC^ (Figure 2B, (Cang et al., 2005; Savier et al., 2017; Tsigankov and Koulakov, 2010, 2006)):

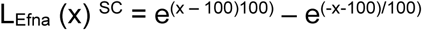

#### Cortical V1 Epha gradients

Estimated gradient of Epha receptors (R _Epha_) in V1 along the medial-lateral axis (x), R _Epha_ (x) ^V1^ (Tsigankov and Koulakov, 2010, 2006) for all genotypes:

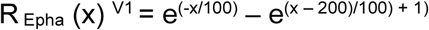

#### Retinal Efna gradients

Measured gradients of Efna ligands in RGCs along the nasal-temporal axis L_Efna_ (x)^retina^ (Savier et al., 2017) for all genotypes :

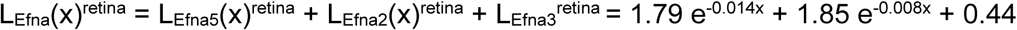

#### Retinal Efna gradients transposition

When transposed along the rostral-caudal axis in the SC (retina -> SC), retinal Efna gradients are flipped along the x axis and become:

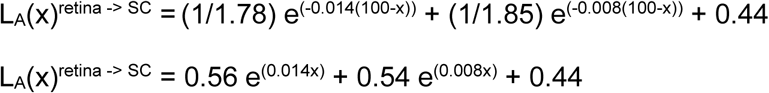

### Intrinsic Dispersion Index (IDI) – Local intrinsic Dispersion variation (IDV) - Alignment Index (AI)

The intrinsic dispersion index and local intrinsic variation, corresponding to the degree of dispersion of the maps are calculated by:

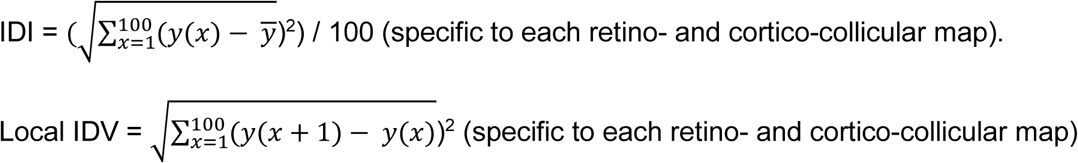

The alignment index is calculated by:

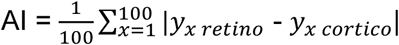

### Statistical analysis

Comparison of relative Efna/ Epha mRNA expression levels in SC and V1 in Isl2-Epha3^KI/KI^ animals relative to WT were performed by one sample t-test, reference value = 1 as described in (Savier et al., 2017). Comparison of relative Efnas expression levels, relative to WT Nasal in Nasal, Central and Temporal WT and Isl2-Epha3^KI/KI^ were performed by the two-way ANOVA without replication (Efna x genotype comparison). Experimental and simulated visual maps were compared using Kolmogorov-Smirnov 2-sample test.

Descriptive analysis of IDV curves were performed using:

- the Jaccard similarity index (J), which indicates the degree of similarity of two sets of data, varies between J = 0 (no similarity) and J = 1 (perfect similarity) and is calculated as follow (Metcalf and Casey, 2016):

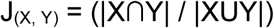
- covariance cov (x, y):

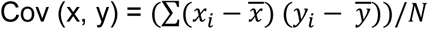

where Cov > 0 indicates similar trend and Cov < 0 indicates opposite trend.

### Data Availability

Leave-one-out ‘Leave-one-out’ script in R: https://github.com/michaelreber/Leave-one-out-LOESS/blob/master/wtloess.R 3-step map alignment model 3-step map alignment script in MATLAB: https://github.com/michaelreber/3-step-Map-Aligment-Model

## Acknowledgements

We are grateful to Amelie Barthelemy and Frank Pfrieger, CNRS UPR3212 Strasbourg France for technical advice on retinal ganglion cell isolation. This work was supported by USIAS – University of Strasbourg, CNRS and start-up funds from the DKJ Eye Institute, Krembil Research Institute, University Health Network.

## Author Contributions

E.S. and M.R. designed and performed the experiments, implemented the 3-step map alignment algorithm, analyzed the results, wrote and edited the manuscript. K.C performed qRT-PCR and projections tracing. J.D. performed algorithm implementation, projections tracing and edited the manuscript. Material requests and correspondence should be addressed to M.R.

## References

Ackman JB, Crair MC. 2014. Role of emergent neural activity in visual map development. Curr Opin Neurobiol 24:166–175. doi: 10.1016/j.conb.2013.11.011

Basso MA, May PJ. 2017. Circuits for Action and Cognition: A View from the Superior Colliculus. Annu Rev Vis Sci 3:197–226. doi: 10.1146/annurev-vision-102016-061234

Bevins N, Lemke G, Reber M. 2011. Genetic dissection of EphA receptor signaling dynamics during retinotopic mapping. J Neurosci Off J Soc Neurosci 31:10302–10310. doi: 10.1523/JNEUROSCI.1652-11.2011

Brown A, Yates PA, Burrola P, Ortuño D, Vaidya A, Jessell TM, Pfaff SL, O’Leary DD, Lemke G. 2000. Topographic mapping from the retina to the midbrain is controlled by relative but not absolute levels of EphA receptor signaling. Cell 102:77–88.

Cang J, Feldheim DA. 2013. Developmental mechanisms of topographic map formation and alignment. Annu Rev Neurosci 36:51–77. doi: 10.1146/annurev-neuro-062012-170341

Cang J, Kaneko M, Yamada J, Woods G, Stryker MP, Feldheim DA. 2005. Ephrin-as guide the formation of functional maps in the visual cortex. Neuron 48:577–589. doi: 10.1016/j.neuron.2005.10.026

Claudepierre T, Koncina E, Pfrieger FW, Bagnard D, Aunis D, Reber M. 2008. Implication of neuropilin 2/semaphorin 3F in retinocollicular map formation. Dev Dyn Off Publ Am Assoc Anat 237:3394–3403. doi: 10.1002/dvdy.21759

Efron B, Tibshirani R. 1991. Statistical data analysis in the computer age. Science 253:390–395. doi: 10.1126/science.253.5018.390

Feldheim DA, Kim YI, Bergemann AD, Frisén J, Barbacid M, Flanagan JG. 2000. Genetic analysis of ephrin-A2 and ephrin-A5 shows their requirement in multiple aspects of retinocollicular mapping. Neuron 25:563–574.

Feldheim DA, Vanderhaeghen P, Hansen MJ, Frisén J, Lu Q, Barbacid M, Flanagan JG. 1998. Topographic guidance labels in a sensory projection to the forebrain. Neuron 21:1303–1313.

Goodhill GJ, Xu J. 2005. The development of retinotectal maps: a review of models based on molecular gradients. Netw Bristol Engl 16:5–34.

Grimbert F, Cang J. 2012. New model of retinocollicular mapping predicts the mechanisms of axonal competition and explains the role of reverse molecular signaling during development. J Neurosci Off J Soc Neurosci 32:9755–9768. doi: 10.1523/JNEUROSCI.6180-11.2012

Hjorth JJJ, Sterratt DC, Cutts CS, Willshaw DJ, Eglen SJ. 2015. Quantitative assessment of computational models for retinotopic map formation. Dev Neurobiol 75:641–666. doi: 10.1002/dneu.22241

Honda H. 2004. Competitive interactions between retinal ganglion axons for tectal targets explain plasticity of retinotectal projection in the servomechanism model of retinotectal mapping. Dev Growth Differ 46:425–437. doi: 10.1111/j.1440-169x.2004.00759.x

Honda H. 2003. Competition between retinal ganglion axons for targets under the servomechanism model explains abnormal retinocollicular projection of Eph receptor-overexpressing or ephrin-lacking mice. J Neurosci Off J Soc Neurosci 23:10368–10377.

Janes PW, Saha N, Barton WA, Kolev MV, Wimmer-Kleikamp SH, Nievergall E, Blobel CP, Himanen J-P, Lackmann M, Nikolov DB. 2005. Adam meets Eph: an ADAM substrate recognition module acts as a molecular switch for ephrin cleavage in trans. Cell 123:291–304. doi: 10.1016/j.cell.2005.08.014

Khachab MY, Bruce LL. 1999. The maturation of corticocollicular neurons in mice. Brain Res Dev Brain Res 112:145–148.

Koulakov AA, Tsigankov DN. 2004. A stochastic model for retinocollicular map development. BMC Neurosci 5:30. doi: 10.1186/1471-2202-5-30

Liang F, Xiong XR, Zingg B, Ji X, Zhang LI, Tao HW. 2015. Sensory Cortical Control of a Visually Induced Arrest Behavior via Corticotectal Projections. Neuron 86:755–767. doi: 10.1016/j.neuron.2015.03.048

Metcalf L, Casey W. 2016. Chapter 2 - Metrics, similarity, and sets In: Metcalf L, Casey W, editors. Cybersecurity and Applied Mathematics. Boston: Syngress. pp. 3–22. doi: 10.1016/B978-0-12-804452-0.00002-6

Owens MT, Feldheim DA, Stryker MP, Triplett JW. 2015. Stochastic Interaction between Neural Activity and Molecular Cues in the Formation of Topographic Maps. Neuron 87:1261–1273. doi: 10.1016/j.neuron.2015.08.030

Peng J, Fabre PJ, Dolique T, Swikert SM, Kermasson L, Shimogori T, Charron F. 2018. Sonic Hedgehog Is a Remotely Produced Cue that Controls Axon Guidance Trans-axonally at a Midline Choice Point. Neuron 97:326-340.e4. doi: 10.1016/j.neuron.2017.12.028

Pfeiffenberger C, Yamada J, Feldheim DA. 2006. Ephrin-As and patterned retinal activity act together in the development of topographic maps in the primary visual system. J Neurosci Off J Soc Neurosci 26:12873–12884. doi: 10.1523/JNEUROSCI.3595-06.2006

Rashid T, Upton AL, Blentic A, Ciossek T, Knöll B, Thompson ID, Drescher U. 2005. Opposing gradients of ephrin-As and EphA7 in the superior colliculus are essential for topographic mapping in the mammalian visual system. Neuron 47:57–69. doi: 10.1016/j.neuron.2005.05.030

Reber M, Burrola P, Lemke G. 2004. A relative signalling model for the formation of a topographic neural map. Nature 431:847–853. doi: 10.1038/nature02957

Rhoades RW, Mooney RD, Fish SE. 1985. Subcortical projections of area 17 in the anophthalmic mouse. Brain Res 349:171–181. doi: 10.1016/0165-3806(85)90141-5

Savier E, Eglen SJ, Bathélémy A, Perraut M, Pfrieger FW, Lemke G, Reber M. 2017. A molecular mechanism for the topographic alignment of convergent neural maps. eLife 6. doi: 10.7554/eLife.20470

Savier E, Reber M. 2018. Visual Maps Development: Reconsidering the Role of Retinal Efnas and Basic Principle of Map Alignment. Front Cell Neurosci 12:77. doi: 10.3389/fncel.2018.00077

Shanks JA, Ito S, Schaevitz L, Yamada J, Chen B, Litke A, Feldheim DA. 2016. Corticothalamic Axons Are Essential for Retinal Ganglion Cell Axon Targeting to the Mouse Dorsal Lateral Geniculate Nucleus. J Neurosci Off J Soc Neurosci 36:5252–5263. doi: 10.1523/JNEUROSCI.4599-15.2016

Simpson HD, Goodhill GJ. 2011. A simple model can unify a broad range of phenomena in retinotectal map development. Biol Cybern 104:9–29. doi: 10.1007/s00422-011-0417-y

Steinmetz CC, Buard I, Claudepierre T, Nägler K, Pfrieger FW. 2006. Regional variations in the glial influence on synapse development in the mouse CNS. J Physiol 577:249–261. doi: 10.1113/jphysiol.2006.117358

Sterratt DC, Hjorth JJJ. 2013. Retinocollicular mapping explained? Vis Neurosci 30:125–128. doi: 10.1017/S0952523813000254

Suetterlin P, Drescher U. 2014. Target-independent ephrina/EphA-mediated axon-axon repulsion as a novel element in retinocollicular mapping. Neuron 84:740–752. doi: 10.1016/j.neuron.2014.09.023

Triplett JW, Owens MT, Yamada J, Lemke G, Cang J, Stryker MP, Feldheim DA. 2009. Retinal input instructs alignment of visual topographic maps. Cell 139:175–185. doi: 10.1016/j.cell.2009.08.028

Triplett JW, Pfeiffenberger C, Yamada J, Stafford BK, Sweeney NT, Litke AM, Sher A, Koulakov AA, Feldheim DA. 2011. Competition is a driving force in topographic mapping. Proc Natl Acad Sci U S A 108:19060–19065. doi: 10.1073/pnas.1102834108

Tsigankov D, Koulakov AA. 2010. Sperry versus Hebb: topographic mapping in Isl2/EphA3 mutant mice. BMC Neurosci 11:155. doi: 10.1186/1471-2202-11-155

Tsigankov DN, Koulakov AA. 2006. A unifying model for activity-dependent and activity-independent mechanisms predicts complete structure of topographic maps in ephrin-A deficient mice. J Comput Neurosci 21:101–114. doi: 10.1007/s10827-006-9575-7

Willshaw D. 2006. Analysis of mouse EphA knockins and knockouts suggests that retinal axons programme target cells to form ordered retinotopic maps. Dev Camb Engl 133:2705–2717. doi: 10.1242/dev.02430

Willshaw DJ, Sterratt DC, Teriakidis A. 2014. Analysis of local and global topographic order in mouse retinocollicular maps. J Neurosci Off J Soc Neurosci 34:1791–1805. doi: 10.1523/JNEUROSCI.5602-12.2014

Zembrzycki A, Stocker AM, Leingärtner A, Sahara S, Chou S-J, Kalatsky V, May SR, Stryker MP, O’Leary DD. 2015. Genetic mechanisms control the linear scaling between related cortical primary and higher order sensory areas. eLife 4. doi: 10.7554/eLife.11416

Zhao X, Liu M, Cang J. 2014. Visual cortex modulates the magnitude but not the selectivity of looming-evoked responses in the superior colliculus of awake mice. Neuron 84:202–213. doi: 10.1016/j.neuron.2014.08.037

